# Black swimming dots in cell culture: the identity, detection method and judging criteria

**DOI:** 10.1101/366906

**Authors:** Xiao Yang, Xinlong Jiang, Wenzhong An, Xiangchen Li, Shuo Wang, Hongtu Li, Zhengchao Wu, Feng Su, Shiliang Ma, Shuang Tang

## Abstract

Black swimming dots (BSDs) — the biological UFO in cell culture — have perplexed biologists for decades. BSDs are extremely tiny dots found in dishes of cultured cells. It is still controversial on their origin and identity. BSDs are very hard to be removed and bring adverse impact to cell experiments (Supplementary Table 1). Here we wish to answer three urgent questions about BSDs. First, is the identity of BSDs nonliving matter or living organism? Second, is there any reliable method to tell whether the donor cattle for FBS production or the animals for primary cells isolation carry BSDs? Third, what are the judging criteria for BSDs when “tiny black dots” were observed? In 2015, we happened to observe BSDs in the perivitelline space of mouse oocytes and embryos; and the cells derived from these mice exhibited typical properties of BSDs infected cells (described in Supplementary Information). With these BSDs infected (BSD+) mice, we demonstrate that BSDs *per se* are nonliving inorganic nanoparticles yet should derive from an unidentified airborne infectious organism. We also suggest that observing the perivitelline space of MII oocytes is an accurate method to detect BSDs in animals. Moreover, we propose some BSDs judging criteria when “tiny black dots” are found in cell or animal samples.

We first found BSDs in mouse embryos in May 2015 (the discovery of BSDs and the phenotypes of BSD+ embryos and cells were detailed described in Supplementary Information; Supplementary Video 4, 5; Extended Data Fig. 1). The *in vitro* cultured zygotes were blocked at 8-cell stage (Extended Data Fig. 1b, c). When examined the zygotes, we surprisingly observed tiny black dots swimming in their perivitelline space. And embryos at each stage also exhibited the same phenomena (Supplementary Video 1). To exclude possible contamination in fertilization, we examined MII oocytes. BSD+ oocytes carried black dots; oppositely, the oocytes from uninfected (BSD-) mice exhibited clean perivitelline space (Supplementary Video 2). The BSD- zygotes normally developed to blastocysts *in vitro* (Extended Data Fig. 1b, c). Moreover, BSD- 8-cell and 32-cell embryos showed clean perivitelline space (Supplementary Video 3). These observations confirmed embryo arrest was induced by BSDs and independent from culture milieu.

To assess the morphology and element composition of BSDs, we performed microscopy and EDX (energy dispersive X-Ray spectroscopy) analysis. Under 400X inverted microscope with phase contrast, BSDs exhibited varying shapes (spheres or rods) and typical features of Brownian motion (Fig. 1a, a’; Supplementary Video 1, 2). Viewed under electronic microscope, BSDs showed tiny size (mainly range from 20~ 500 nm), oval-shaped morphology (Fig. 1b-d). The EDX spectra showed high-intensity peaks of Ca, P, O and C (Fig. 1e). The Ca/P/O/C atomic ratio implied BSDs might be inorganic nanoparticles composed of HAP (hydroxyapatite) and CaCO_3_ (Fig. 1g). It was reminiscent of previous papers on nanobacteria (NB) ^1–3^.

**Figure 1.**
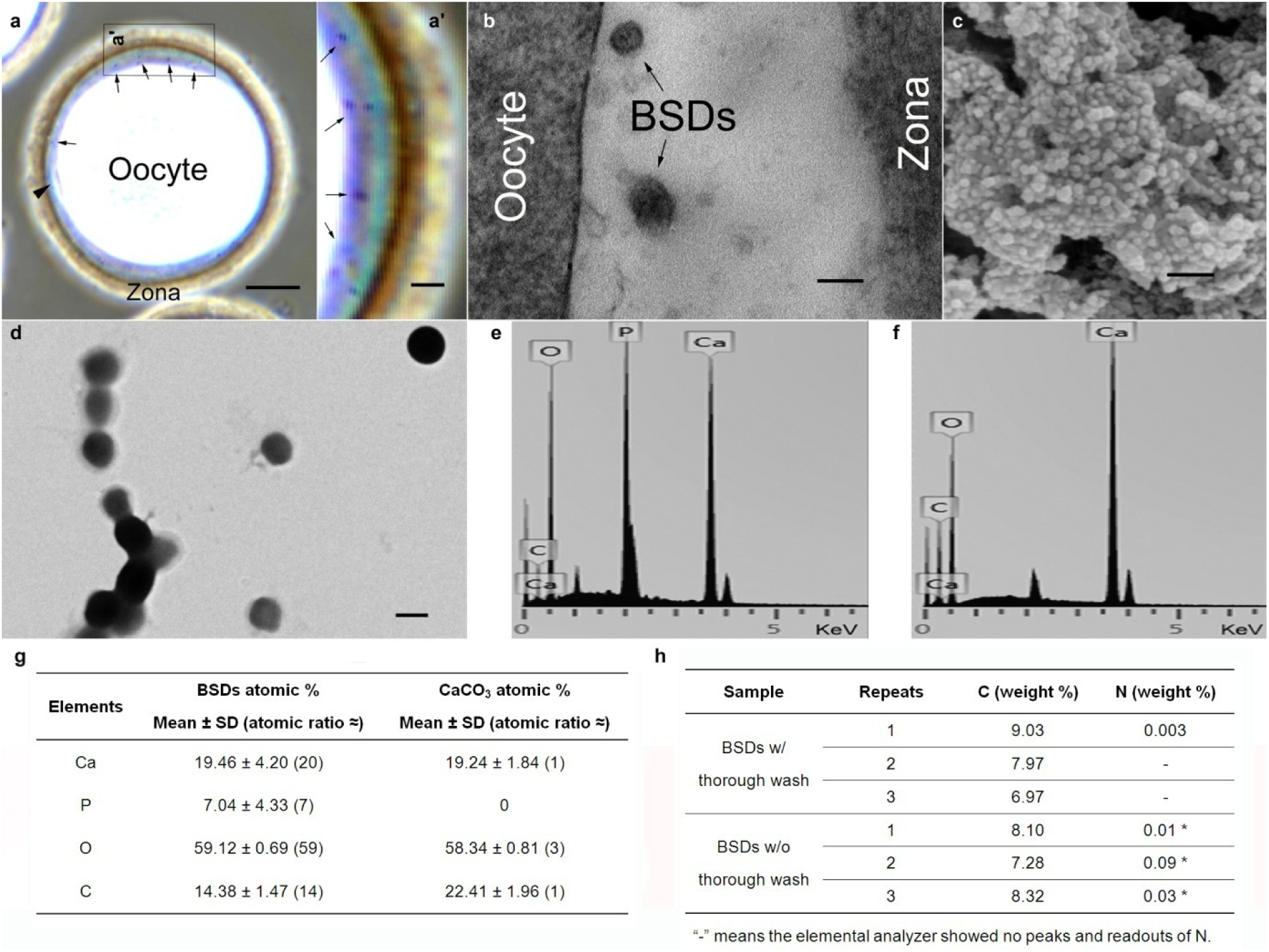
The morphology and element composition of BSDs. **a**, Phase contrast image of BSDs in the perivitelline space of a mouse oocyte. Arrows indicated BSDs; arrowhead indicated polar body. Scale bar = 10 μm. **a’**, The enlargement of the boxed region in **a**. Scale bar = 2 μm. **b**, TEM image of BSDs in the perivitelline space of a mouse oocyte. Scale bar = 100 nm. **c**, SEM image of BSDs collected by centrifugation from BSD+ cell culture. Scale bar = 100 nm. **d**, Negative staining of large BSDs observed under TEM. Scale bar = 500 nm. **e, f**, Representative EDX spectra of BSDs (**e**) and CaCO_3_ (**f**). **g**, EDX data. Mean ± SD was calculated from the data in three independently repeated experiments. CaCO_3_ was used as EDX reference and its atomic ratio was approximately equal to 1/3/1 for Ca/O/C. The Ca/P/O/C atomic ratio of BSDs roughly matched the ratio of HAP (molecular formula: Ca_10_(PO_4_)_6_(OH)_2_) and CaCO_3_. **h**, The data of C/N elemental analysis. “–” No peaks and readouts in the elemental analyzer. **\*** Interestingly, low amount of N could be detected in BSDs without thorough wash. That should come from albumin or other proteins absorbed on BSDs in cell media, as previously reported ^3,8,9^. The sample treatments were as described in the “Electronic microscopy and EDX” and “Nitrogen content in BSDs” sections of the **Methods**.

Concerning BSDs’ identity, NB seemed to be closest to the truth since the characteristics of NB and BSDs were quite similar (Fig. 1) (Posts 36, 37, 39, 41 in Supplementary Table 1) ^1,4^. The other speculations (Supplementary Table 1), such as mycoplasma, bacteria, Achromobacter ^5^ and cell debris, were easy to be experimentally excluded (described in Supplementary Information; Supplementary Video 6; Extended Data Fig. 4, 5, 6; Extended Data Table 1). Here we emphatically discuss why BSDs are not NB, and then present our understandings on BSDs’ identity based on our empirical evidence.

Folk first proposed the concept “nanobacteria” and their possible biogenic mineral deposition function ^6^, then Kajander group declared finding of NB in blood and commercial FBS, and called NB as “biofilm producing ORGANISMS” ^7^. Subsequently, they published their most controversial paper in which NB caused nephrolith as the apatite-producing pathogen and were described as the smallest bacteria with biotic properties such as containing DNA and protein, self-replication, sensitive to antibiotics, growth on Loeffler medium ^1^. Biomineralization is closely related to human health ^8–11^. If NB were real, many unsolved biological problems, e.g. BSDs or lithiasis, would be well answered ^8,9,12^. However, NB were questioned for lack of solid evidence ^8,9,12^ and other independent groups stated NB were actually HAP ^2^ or CaCO_3_ ^3^ nanoparticles. Likewise, our analysis also showed BSDs *per se* were merely nonliving inorganic nanoparticles, not Kajander’s NB, for two reasons:

I. EDX showed BSDs were nanoparticles comprising Ca/P/O/C. If BSDs were NB and contained nucleic acids or protein ^1^, at least they should show a certain proportion of N since EDX probing volume is 1 μm^3^, greater than BSDs volume ^13,14^. However, all groups obtained consistent results without detection of N element (Fig. 1e, g) ^1–3^. To further verify that, we determined N content in BSDs by elemental analyzer and obtained negative results (Fig. 1h). We also tried to detect DNA using Kajander’s method ^1^, yet, similar to previous reports ^2,3^, found no DNA.

II. Kajander group reported aminoglycoside and tetracycline prevent NB multiplication ^1,15^. We tested the antibiotics, but they did not work (Extended Data Table 1; Extended Data Fig. 4). We also attempted to cultivate BSDs on Loeffler medium ^1^, but nothing grew out as Cisar et al. reported ^2^.

Therefore, the existing BSDs could not be interpreted as NB. However, the verdict that BSDs are nonliving inorganic nanoparticles has a significant defect: it cannot explain biotic properties of BSDs.

First of all, BSDs demonstrated airborne infectivity in both animals and cultured cells. BSD-mice were infected with BSDs when they were raised in the room of BSD+ mice even without direct contact (Fig. 2a). So did BSD- cells in the same incubator with BSD+ cells (Fig. 2a) (consistent with Posts 30, 34 in Supplementary Table 1). BSDs could also be transmitted via contact infection. Serum from BSD+ mice or media from BSD+ cells could cause BSDs infection when added into dishes of BSD-cells. Notably, they lost infectivity after 134°C, 4 h autoclave (Fig. 2b). Moreover, after BSD- mice and cells were infected, they could serve as new source of infection to cause BSDs in other BSD- mice and cells (Fig. 2c). The above observations conform to Koch’s postulates and the only way to explain BSDs’ infectivity is that they should derive from an unidentified pathogen because only in organisms can INFECTIVITY exist, but not in any inorganics.

**Figure 2.**
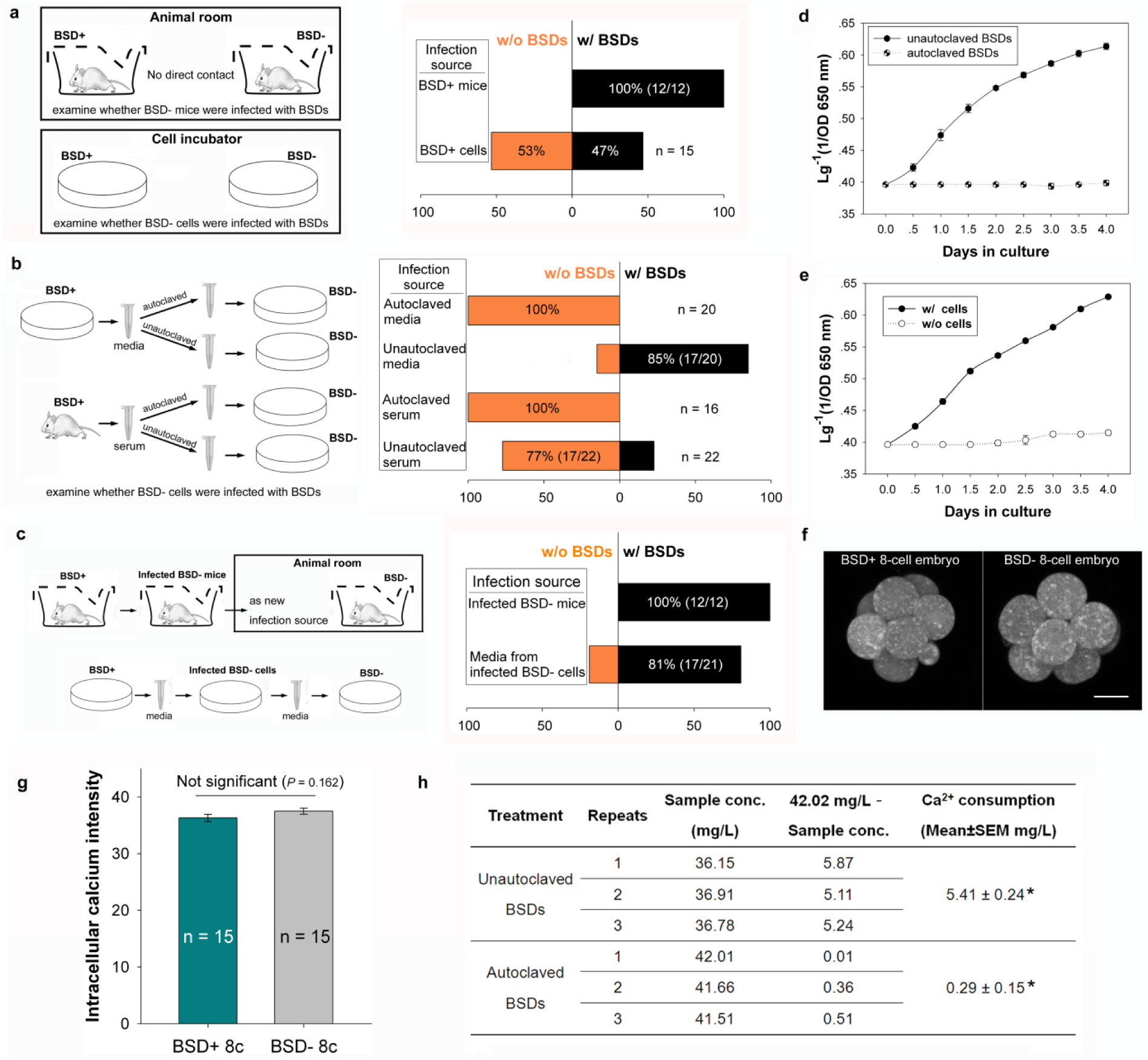
The biotic properties of BSDs. **a**, The airborne infectivity of BSDs. BSDs could be transmitted from BSD+ mice or cells to BSD- ones via air. **b**, The contact infectivity of BSDs. BSD- cells were infected with BSDs by addition of BSD+ serum or media. In contrast, BSD+ serum or media lost infectivity after autoclave. **c**, The BSD-mice and cells acquired the same infectivity after they were infected with BSDs. **d**, Growth curves of unautoclaved and autoclaved BSDs showing BSDs *per se* did not possess the infectivity. Each bar represented SEM calculated from three independently repeated experiments. **e**, Growth curves of BSDs in dishes with or without cells showing BSDs proliferation was dependent on cells. Each bar represented SEM. **f**, Representative image of the intracellular calcium in BSD+ and BSD- 8-cell embryos. Scale bar = 20 μm. **g**, The intracellular calcium intensity of 8-cell embryos. Each column denoted mean ± SEM. Difference between BSD+ and BSD-groups were analysed by Student’s *t*-test. The data passed the Normality Test (*P* = 0.790) and Equal Variance Test (*P* = 0.689). The *t*-value was −1.436 with 28 degrees of freedom. **h**, The AAS data of calcium concentration in culture media. Unautoclaved or autoclaved BSDs were added into the cell dishes. At the 4^th^ day, media were collected and analyzed by AAS. Difference between two groups were analysed by Student’s *t*-test. The data passed the Normality Test (*P* = 0.588) and Equal Variance Test (*P* = 0.677). The *t*-value was 18.425 with 4 degrees of freedom. **\*** There is a statistically significant difference between two groups (*P* ≤ 0.001). The preparation and autoclave treatment of media, serum and BSDs were as described in the “Filtration and autoclave treatment” and “Mouse serum preparation and autoclave treatment” sections of the **Methods**.

What further supporting this point is that the infectivity did not lie in BSDs *per se*. When inoculated into cultured cells, unautoclaved BSDs multiplied quickly, yet autoclaved BSDs lost their infectivity (Fig. 2d). Besides, even if BSDs were removed, media from BSD+ cells could cause BSDs in BSD- cells (Fig. 2b). It reinforced BSDs *per se*, as inorganic nanoparticles, were impossible to be infectious.

Next, BSDs proliferation was dependent on cells. When media from BSD+ cells were added into dishes without cells or with cells, the multiplication displayed two distinct patterns. BSDs multiplied rapidly when cells existed; in contrast, BSDs hardly proliferated without cells (Fig. 2e).

Finally, BSDs (or rather, the unidentified pathogen of BSDs) might affect calcium metabolism, but not intracellular calcium concentration, of the cells. The compaction failure (Extended Data Fig. 1b, c) reminded us to check the intracellular calcium of 8-cell embryos. Unexpectedly, BSD+ 8-cell embryos showed no difference in calcium intensity from BSD-ones (Fig. 2f, g). These results prompted us to speculate whether the unidentified pathogen caused calcium influx into the cells and then the cells produced BSDs for detoxification of overhigh intracellular calcium ^16–18^. The AAS (atomic absorption spectroscopy) data supported this speculation. When BSDs multiplied, calcium in the culture media were depleted greatly (5.41 ± 0.24 mg/L, Fig. 2h). In comparison, autoclaved BSDs hardly multiplied, and calcium concentration of the media dropped by only 0.29 ± 0.15 mg/L (Fig. 2h). These results were consistent with the facts that no report showed the nucleation and growth of inorganic apatite under physiologic conditions ^15^.

The above properties exclusively belong to living organisms. Taken together, we propose BSDs *per se* are nonliving inorganic nanoparticles yet should derive from an unidentified pathogen which should have the following characteristics: airborne transmitted; cell-dependent; insensitive to antibiotics; filterable through 0.1 μm membrane. Furthermore, it may accelerate calcium metabolism of the cells.

Distinguishment of BSD+ animals is crucial for serum industry and the credibility/reproducibility of mechanism research using cells or animals. During our investigation, we noticed BSDs were seen in all BSD+ embryos, but not in all BSD+ cell samples. To assess whether observing perivitelline space of MII oocytes is a reliable method to inspect BSDs infection, we collected MII oocytes and corresponding MTFs (mouse tail-tip fibroblasts) from 9 BSD- and 47 BSD+ mice, respectively. The oocytes and MTFs from BSD- groups were clear of BSDs. In contrast, all oocytes from BSD+ group, without exception, showed BSDs swimming in their perivitelline space. However, BSDs were seen in only 12 out of 47 MTFs from BSD+ mice (Extended Data Fig. 2). In addition, we did not observe BSDs in serum from BSD+ mice although the serum possessed infectivity. Therefore, no BSDs in cells and serum did not mean they were not infected and this could also explain why we faced so many cell lines with BSDs (Supplementary Table 1). Accordingly, observing perivitelline space of MII oocytes could tell BSDs infection accurately. In further studies, oocytes from other species should be checked to see if our proposed method was feasible. If validated, this method might be added into the pathogen inspection list (especially for cattle and laboratory animals).

Due to no criteria, although many laboratories met BSDs, few articles have been published and scientists had to discuss “cells with black dots” on online bioforums (Supplementary Table 1). Based on our investigation and bioforums, we propose some judging criteria for BSDs (Extended Data Fig. 3c). I. BSDs demonstrate Ca, P, O, C peaks in EDX spectra and no N in elemental analysis. II. BSDs are highly infectious. III. BSDs proliferation is dependent on cells. IV. BSDs keep Brownian motion and present nano-sized spherical or rodlike morphology. V. BSDs do not cause pH change and turbidity of culture media. VI. BSDs cannot be cultivated on agar plates or broths. VII. BSDs are resistant to antibiotics. Of note, some of “tiny black dots”, even if they are very like BSDs in appearance ^5,8^, may not be BSDs. Sometimes nanoscale biominerals might exist in normal organisms ^8,10,19^, so only the black dots meeting the above criteria could be identified as BSDs.

Here, we demonstrate BSDs possess obvious biotic-abiotic duality. However, previous studies concentrated much attention on “dots” yet neglected their links with cells. Kajander group observed biotic properties of NB, yet regrettably, their explanation to NB was too subjective ^12^. Cisar et al. and Young group only focused on the dots, so they obtained correct but biased conclusion, namely, inorganic nanoparticles ^2,3,8,19^. Therefore, Ciftcioglu and McKay argued nonbiogenic crystallization cannot explain the biogenic-like properties of NB ^15^. But anyway, these studies provided reference for our investigation and also supported our results from a certain aspect. Based on our results of BSDs’ identity, we suggest a new direction that future investigations should focus on both the cells and the dots. The right direction is decisive for scientific research, otherwise, only can we reach a specious and incomplete conclusion (Extended Data Fig. 3a). In addition, the urgent problem of now is that we faced many BSDs infected FBS and cells. We put forward a simple method to screen out BSD+ animals (Extended Data Fig. 3b). Before the final solution of BSDs problem, this method has great scientific and economic benefits to avoid BSDs. Furthermore, with BSDs’ judging criteria, systematic investigations will gradually accumulate and replace the groundless speculations on BSDs. The imperfection of this study is that we have not established a pure culture. This conundrum requires a multidisciplinary cooperation (e.g. microbiology, embryology, cytology, virology etc.) for final identification of BSDs’ pathogen.

**Supplementary Information** The files contain Supplementary Text and Data; Supplementary Table 1; Supplementary Video 1–6.

- **Supplementary Video 1** | **BSDs in the perivitelline space of BSD+ embryos at each stage.** The embryos were recorded immediately after collection. Zy, zygote; 2c, 2-cell embryo; 4c, 4-cell embryo; 8c, 8-cell embryo; c8c, compacted 8-cell embryo; 16c, 16-cell morula; 32c, 32-cell morula; Bl, blastocyst. The boxed region was zoomed in to show BSDs more clearly.

- **Supplementary Video 2** | **BSDs in the perivitelline space of BSD+ oocytes vs. no BSDs in the perivitelline space of BSD-oocytes (BSD- control).** The boxed region was zoomed in to show BSDs more clearly in the left video. The z-axis focus was adjusted to show no BSDs in different layers of BSD- oocytes in the right video.

- **Supplementary Video 3** | **The 8-cell and 32-cell embryos from BSD- mice showing clean perivitelline space (BSD- control).**

- **Supplementary Video 4** | **BSDs in the dish of cells isolated from BSD+ mice (a) vs. no BSDs in the dish of cells isolated from BSD- mice (b).**

- **Supplementary Video 5** | **BSDs proliferation in dishes of active-state cell strains.** Even if BSDs were washed off at the day before (**a**), they would grow full of the dish overnight (**b**). The proliferation of BSDs after another day of culture (**c**).

- **Supplementary Video 6** | **The morphological difference between BSDs (a) and Achromobacter (b).**

## Acknowledgements

We thank Dexian Zhang for assistance in animal management; Lixin Yin for SEM and EDX analysis; Zhengzheng Ma for AAS analysis. This research was supported by the National Natural Science Foundation of China (Grant 31301201 to S.T.).

## Author Contributions

S.T. conceived the idea and designed the research; X.Y., X.J., W.A., S.W., Z.W & S.T. performed the experiments; X.L., H.L. & F.S. provided some materials and discussed the results; X.Y., X.J., W.A., H.L., S.M. & S.T. analyzed the data; S.T. wrote the manuscript.

## Author Information

The authors declare no competing financial interests. Readers are welcome to comment on the online version of the paper. Correspondence and requests for materials should be addressed to S.T.

## Methods

### Reagents

All reagents were purchased from Sigma unless otherwise stated. DMEM/F-12 and FBS (fetal bovine serum) were from Gibco.

### Animals

We declare our respect to the welfare of laboratory animals in the experiments. SPF (Specific Pathogen Free) ICR mice were purchased from Jinan Pengyue Lab animals Co. Ltd. (Jinan, Shandong, China), Liaoning Changsheng Biotechnology Co. Ltd. (Liaoning, China) or Beijing Huafukang Bioscience Co. Inc. (Beijing, China). The mice were accordingly named as JN, LN and BJ, respectively (Extended data Fig. 1a, b). The mice were maintained with food and water *ad libitum* on a 12 h light/dark cycle under controlled temperature (23 to 25°C) in the Lab Animal Center of Shenyang Agricultural University. Since BSDs were infectious (Fig. 2a), the mice from different Company were raised in independent rooms. The 6 to 8-week-old female mice and 12 to 24-week-old male mice were used for the experiments. All protocols were approved by the Institutional Animal Care and Use Committee of Shenyang Agricultural University.

### Designation of BSD- (healthy, no BSDs) and BSD+ (BSDs infected) mice and cells

The mice (JN and LN-b) showing clean perivitelline space and normal *in vitro* development were designated as BSD- (uninfected mice). The mice (LN-a and BJ) showing BSDs in perivitelline space and 8-cell arrest *in vitro* were designated as BSD+ (infected mice) (Extended data Fig. 1a, b). The JN (BSD-) and LN-a (BSD+) mice were used in most part of this study. The cells from BSD- mice were designated as BSD- cells, so were BSD+ cells.

### Collection of embryos and *in vitro* culture

The medium for collection of oocytes and embryos was the modified potassium simplex optimized medium containing 20 mM HEPES, 4 mM NaHCO_3_ and 1 mg/mL BSA (HKSOM) or M2 (M7167, Sigma). The culture medium was the modified potassium simplex optimized medium with essential and non-essential amino acids containing 8 mg/mL BSA (KSOM) or M16 (M7292, Sigma); in serum group, they were supplemented with additional 10% FBS.

To collect MII oocytes, female mice were superovulated with 10 IU of PMSG (pregnant mare serum gonadotropin), followed 48 h later by 10 IU of HCG (human chorionic gonadotropin). The oviducts were dissected from superovulated females at 16 h post-HCG. Cumulus-oocyte complexes (COCs) were released by teasing apart the ampulla of oviducts. The cumulus cells were dispersed by treatment with 1 mg/mL hyaluronidase in PBS.

To collect embryos at each stage, the superovulated female mice were caged with male mice. Mating was confirmed by identification of a vaginal plug on the next morning. The zygotes were recovered from oviducts at 16 h post-HCG. The cumulus cells were dispersed by treatment with 1 mg/mL hyaluronidase in PBS. The 2-cell, 4-cell, 8-cell, compacted 8-cell and 16/32-cell (morula) embryos were recovered from females by teasing the excised oviducts into pieces at 40 h, 54 h, 64 h, 68 h, 86 h post-HCG, respectively. Blastocysts were obtained by flashing dissected uteri at 98 h post-HCG.

As for *in vitro* culture, every 8~12 zygotes were cultured in a 40 μl droplet of medium at 37.5°C under a humidified atmosphere of 5% CO_2_ in air (this time point was defined as ‘0 h’ in the present study). The development to 8-cell, compact 8-cell, morula and blastocyst stage was examined at 48 h, 54 h, 72 h and 84 h, respectively.

### *In vivo* development of embryos

Female mice were mated with fertile males to induce pregnancy (vaginal plug = day 1 of pregnancy). Parturition events were monitored from days 17 through 21 by observing mice daily in the morning and evening.

### Cell culture

The cell media were DMEM/F-12 containing 10% FBS. The establishment and manipulation of MEFs (mouse embryonic fibroblasts) and MTFs (mouse tail-tip fibroblasts) were performed according to the previous papers ^20,21^. Since BSDs are infectious (Fig. 2a), BSD- and BSD+ cells were maintained in different cell culture rooms. In BSDs’ infectivity experiment, unautoclaved or autoclaved media (or serum) were added into cell media in 1: 10 dilution. The unautoclaved or autoclaved BSDs were supplemented at an OD_650nm_≈0.003 in cell media.

### Filtration and autoclave treatment

To collect autoclaved media, cell media containing BSDs were autoclaved at 134 °C for 4 h, then centrifuged at 21,000 g for 15 min. The supernatant was filtered through a 0.1 μm pore size membrane (autoclaved media). To collect autoclaved BSDs, cell media containing BSDs were centrifuged at 21,000 g for 15 min. The precipitates were washed three times with ddH_2_O and then autoclaved at 134 °C for 4 h. The precipitates were used by resuspension in cell media at an OD650nm≈0.003 (autoclaved BSDs). For unautoclaved media or BSDs, the procedure of 134°C for 4 h was omitted.

### Mouse serum preparation and autoclave treatment

After coagulation, whole blood was centrifuged at 5,000 g for 10 min. The supernatant was collected and filtered through a 0.1 μm pore size membrane. For autoclaved serum, same protocols as above were utilized.

### BSDs observation under inverted microscope

The embryos or cells were observed and captured under a Nikon Ti-S inverted microscope using 40X objective with Ph2 phase contrast ring. The hypertonic HKSOM (the line of volumetric flask was moved down about 0.2% of total volume; e.g. to prepare 500 mL HKSOM, the final volume will be 499 mL) was used for clear observation of the perivitelline space.

### Mycoplasma PCR detection

Mycoplasma was detected using LookOut Mycoplasma PCR Detection Kit (Sigma) according to user’s manual. The negative samples should show a band at approximately 481 bp and mycoplasma positive samples would show bands in the range of 270 ± 8 bp. The oocytes, MTFs and blood from BSD+ and BSD- mice were examined. Blood DNA was isolated with a Wizard Genomic DNA Purification Kit (Promega).

### Antibiotic treatment

For BSD+ embryo *in vitro* culture, the antibiotics were respectively added into KSOM in following concentrations: penicillin/streptomycin 100 U/mL/100 μg/mL, amphotericin 2.5 μg/mL, ciprofloxacin/piperacillin 10 μg/mL each ^5^, gentamicin 50 μg/mL, tetracycline 10 μg/mL, minocycline 10 μg/mL. The *in vitro* development was monitored. In control group, BSD- zygotes were cultured in KSOM without antibiotics. We also tried 2 and 5-fold of routine doses, yet they did not work either (moreover, some antibiotics exerted harmful influence on embryos in high doses).

For cell culture, the antibiotics were added into cell media and PBS. BSDs in BSD+ cell dishes were washed off with PBS, and then media with antibiotics were changed into the dishes. The cells were observed to examine the effects of antibiotics on BSDs.

### Achromobacter inoculation and antibiotic treatment

Achromobacter *xylosoxidans* (ACCC10298, Agricultural Culture Collection of China) were inoculated into BSD- cells. After Achromobacter growing out, media containing 10 μg/mL ciprofloxacin/piperacillin were changed into the Achromobacter inoculated or BSD+ cell dishes, respectively. In the first few days, media containing ciprofloxacin/piperacillin were changed from time to time according to the growing state of Achromobacter or BSDs. Then exchange of media was performed every three days. The growth of Achromobacter and BSDs was monitored.

### Electronic microscopy and EDX (energy dispersive X-Ray spectroscopy)

For BSDs in embryos, sample preparation was performed according to routine protocols with the following modifications. The embryos were fixed in 2.5% glutaraldehyde in PBS for 3 h at 4°C. The embryos were repeatedly washed three times in PBS for 1 h. Then they were thoroughly washed three times in saline for 1 h. Low melting point agarose (LMP) was prepared in a 3 mm^3^ cubic mold. After solidification, a 1 mm diameter screwdriver head was passed back and forth on the flame; stopped for a short while; and then pressed down vertically into the LMP agarose block to form a 1 mm^3^ microwell. The embryos were transferred into the microwell and covered with 37°C LMP. This agarose block was then dehydrated, embedded and ultramicrotomed. Sections were placed on a copper grid and directly observed under a Hitachi HT7700 TEM (transmission electronic microscope).

For BSDs in cell culture, BSDs were collected by centrifugation at 21,000 g for 15 min. The precipitates were thoroughly washed three times with ddH_2_O and then dried. The dried precipitates were adhered on a carbon-tape and sprayed with golden particles and then analyzed under a Zeiss Ultraplus SEM (scanning electronic microscope) with an X-Max^N^ EDX device (Oxford Instruments). To avoid carbon-tape C interference, we chose thick precipitates for EDX analysis.

To observe large BSDs (up to 500 nm), a copper grid was dipped into resuspended BSDs in ddH_2_O and stained in phosphotungstic acid, and then observed by HT7700 TEM.

### Calcium detection in 8-cell embryos

The calcium in 8-cell embryos from BSD+ and BSD- mice were detected with Fluo3-AM (Solarbio, Beijing, China) according to user’s manual. Samples were detected at 488 nm under a Nikon A1 laser scanning confocal microscope. The intracellular calcium intensity was analyzed by using ImageJ software according to the User Guide (https://imagej.nih.gov/ij/).

### Calcium consumption of cell culture media

Unautoclaved or autoclaved BSDs were added into BSD- cells. After 4 days of culture, media were collected and centrifuged at 21,000 g for 15 min. The supernatant was filtered through a 0.1 μm pore size membrane. The Ca^2+^ concentration of media were measured with PinAAcle 900F atomic absorption spectrometer (PerkinElmer). DMEM/F-12 contain 42.02 mg/L Ca^2+^, so Ca^2+^ consumption was equal to (42.02 mg/L – measured concentration).

### Nitrogen content in BSDs

BSDs were collected by centrifugation at 21,000 g for 15 min and the precipitates were thoroughly washed three times with ddH_2_O and then dried. The element contents of N and C in BSDs were determined by Vario EL elemental analyzer (Elementar) or EURO EA elemental analyzer (EURO Vector).

Since Martel et al. reported that serum albumin might be associated to inorganic nanoparticles ^3,8^, BSDs without thorough wash (the BSDs precipitates were resuspended in ddH_2_O, then spinned down and dried) were also examined.

### Statistics

No statistical methods were used to predetermine sample size. No randomization method was used. The investigators were not blinded during experiments.

Each experiment was independently performed at least three times. The exact values of sample number (n) were shown in the figures. The data were analyzed by Student’s *t*-test or one-way ANOVA using Sigmastat3.5 (Systat Software Inc.). Differences between experimental groups were considered significant at *P* < 0.05. The exact values for both significant and non-significant *P* values were provided in the figures or figure legends. The *F* values and degrees of freedom for one-way ANOVA and *t*-values and degrees of freedom for Student’s *t*-test were provided in the figure legends.

---

**Extended Data Figure 1.**
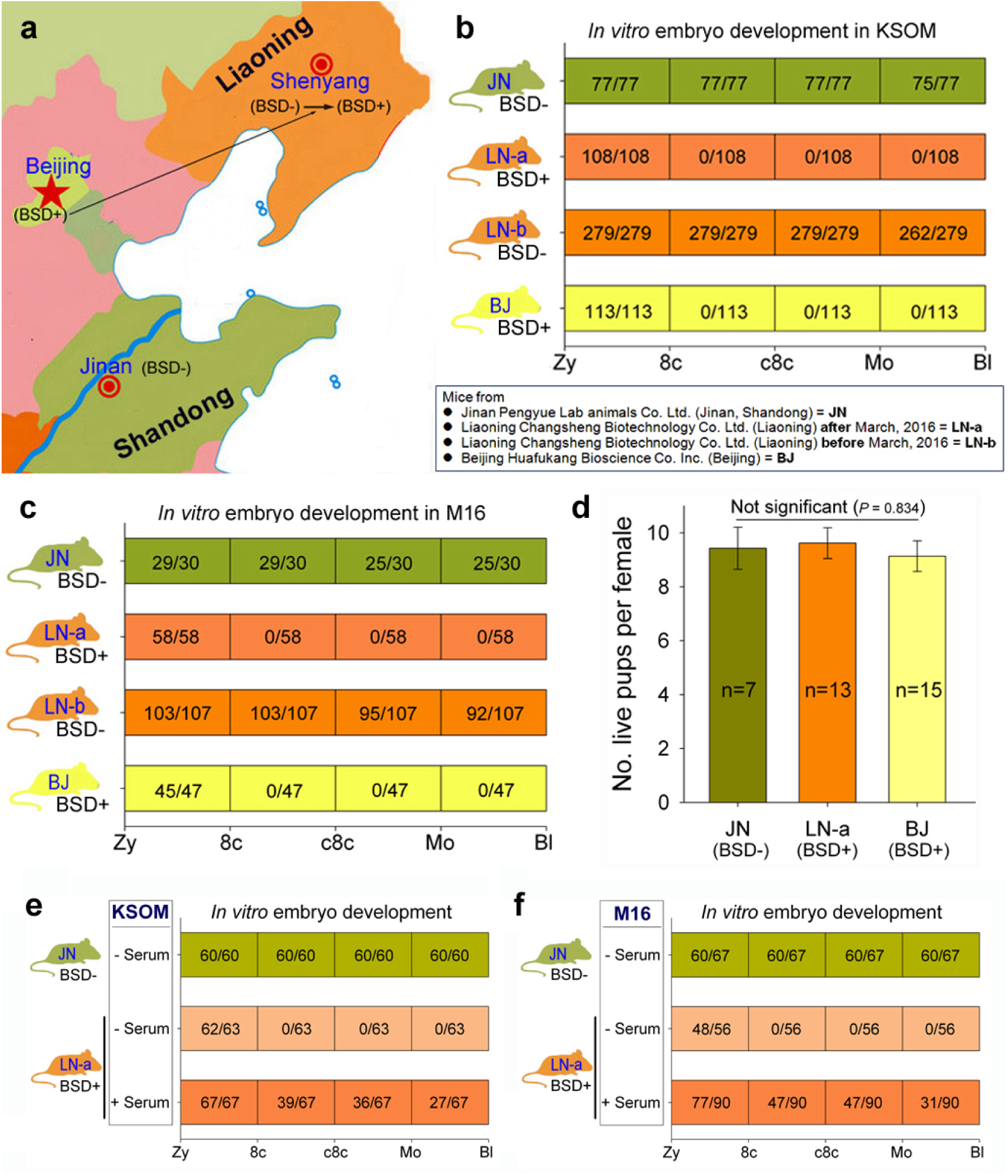
Discovery of BSDs in mouse preimplantation embryos. **a**, The map showing the regions where the different Lab Animal Company located. In Beijing, we first observed BSDs infected mice (BSD+). After these mice were introduced from Beijing to Liaoning province (long arrow), the healthy mice (BSD-) were infected with BSDs (short arrow). The mice from Jinan were clear of BSDs (BSD-). **b**, *In vitro* development of BSD+ and BSD- embryos in KSOM. Designation of JN, LN-a, LN-b, BJ mice and BSD-, BSD+ mice were as described in the boxed text and the “Animals” and “Designation of BSD- (healthy, no BSDs) and BSD+ (BSDs infected) mice and cells” sections of the **Methods**. **c**, *In vitro* development of BSD+ and BSD- embryos in M16. **d**, *In vivo* development of BSD+ and BSD- embryos. The parturition time was at day 20 or 21. Each column denoted mean ± SEM of the live pups number per female. Differences among groups were analysed by One-Way ANOVA. The data passed the Normality Test (*P* = 0.428) and Equal Variance Test (*P* = 0.911). The *F* value was 0.182 with 32 degrees of freedom. **e, f**, The effects of serum supplementation on the *in vitro* development of BSD+ and BSD- embryos cultured in KSOM (**e**) or M16 (**f**).

**Extended Data Figure 2.**
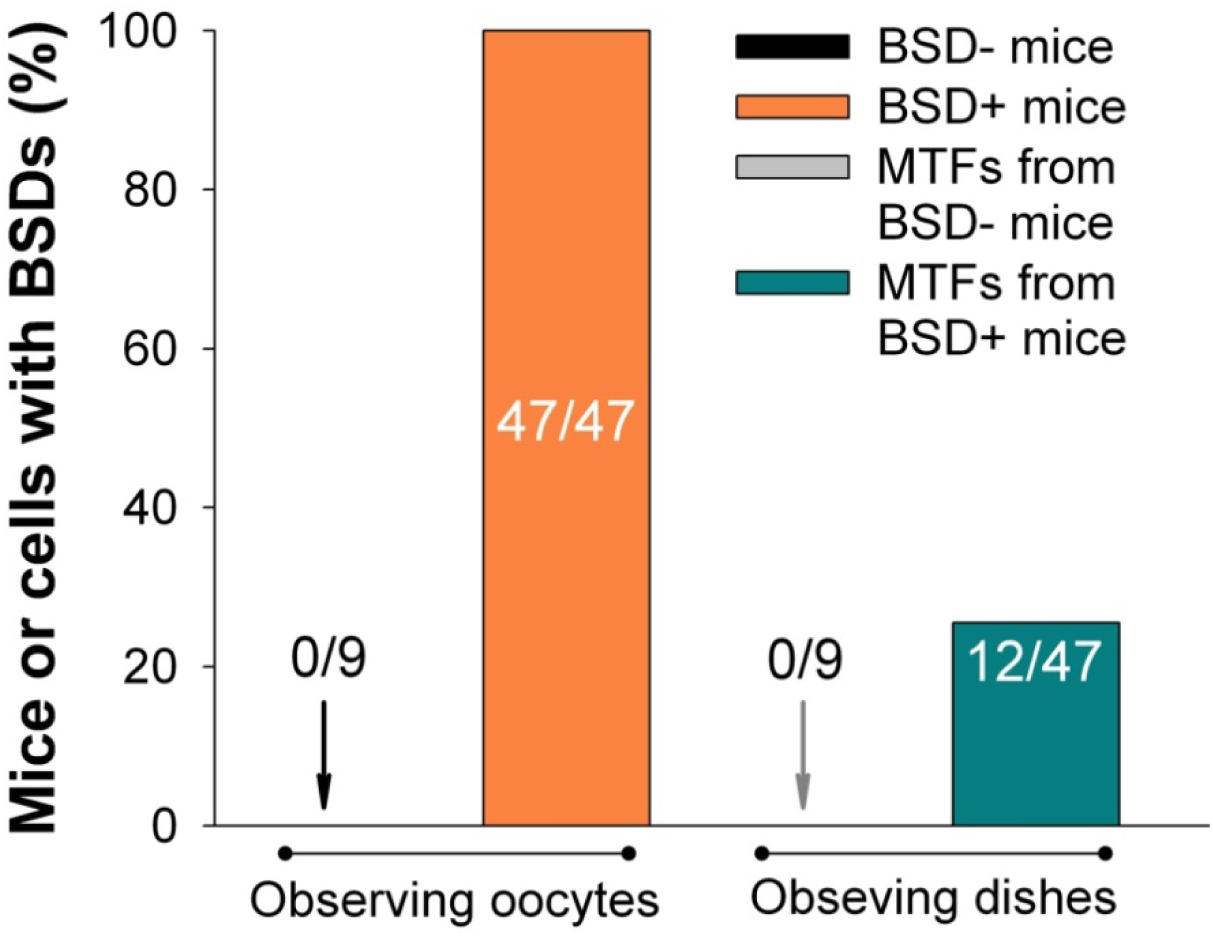
Detection of BSDs infection in animals. MII oocytes and corresponding MTFs isolated from 9 BSD- and 47 BSD+ mice were examined under inverted microscope. Each column denoted the percentage of the mice (determined by observing the perivitelline space of oocytes) or MTFs (determined by observing the cell dishes) with BSDs.

**Extended Data Figure 3.**
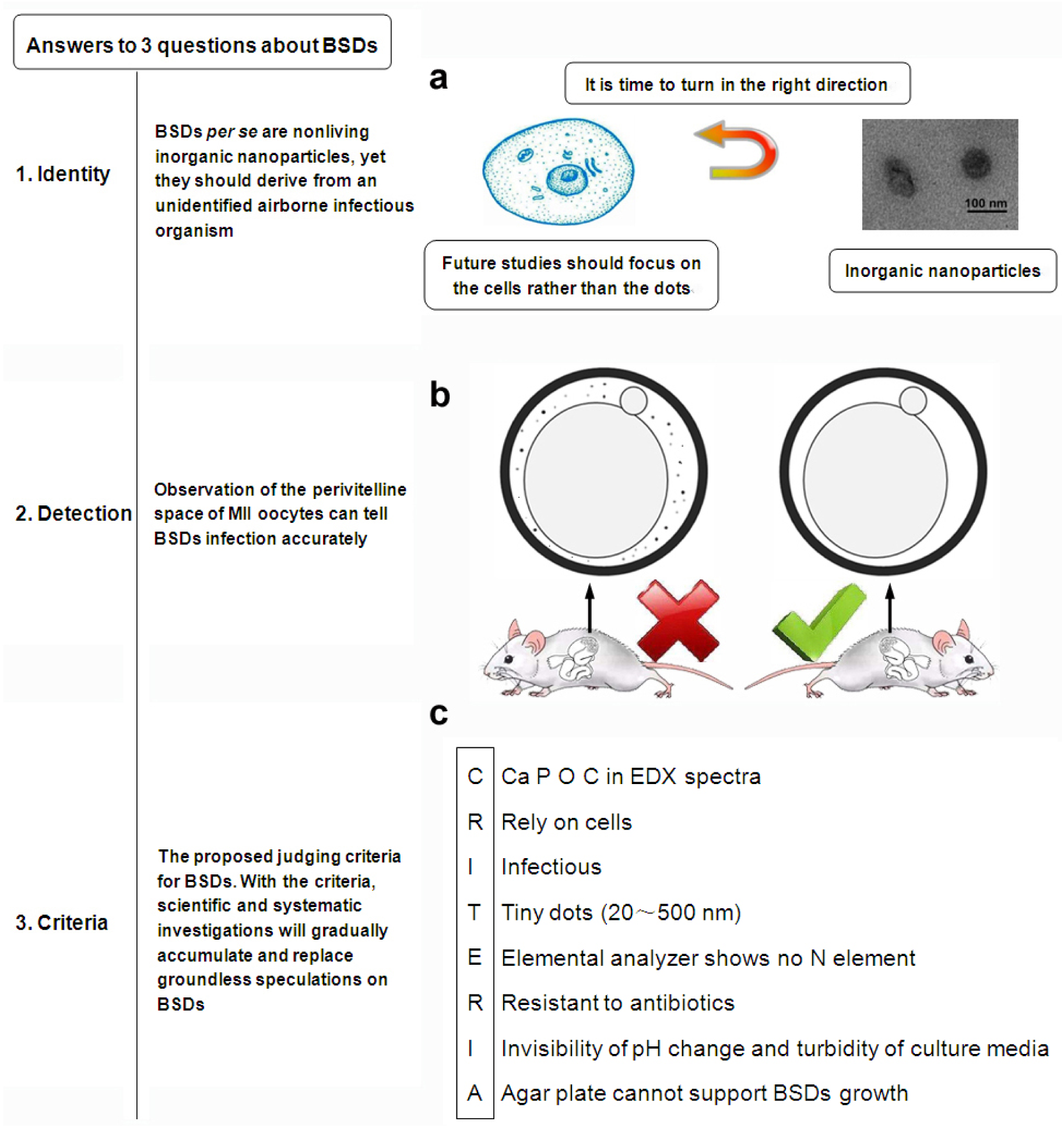
Schematic diagram. The answers to three major questions about BSDs’ identity (**a**), detection method (**b**) and judging criteria (**c**).

**Extended Data Figure 4.**
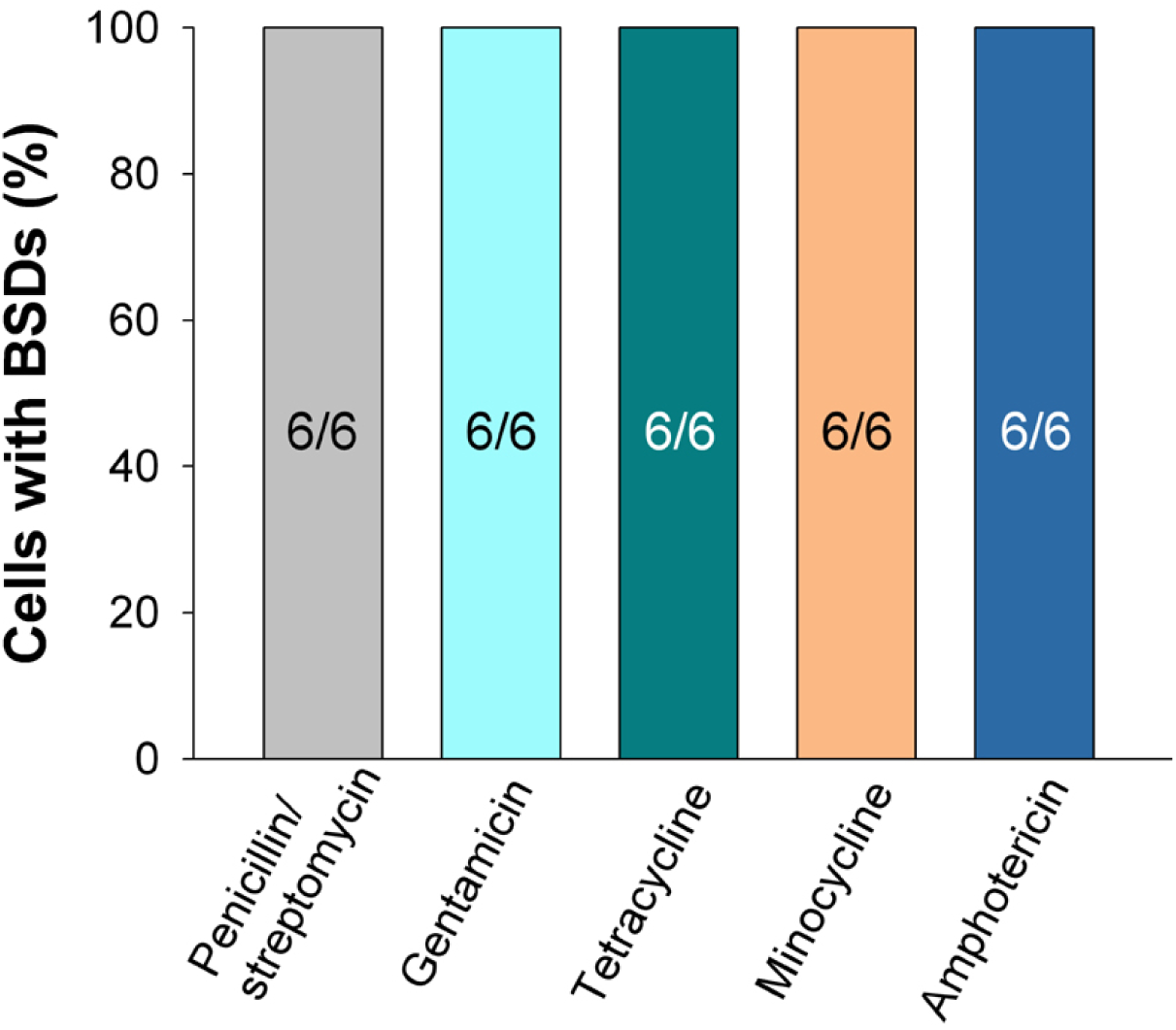
The effects of antibiotics on BSDs in cell culture. BSD+ cells were cultured in media containing the indicated antibiotics. Each column denoted the percentage of the cultured cells with BSDs.

**Extended Data Figure 5.**
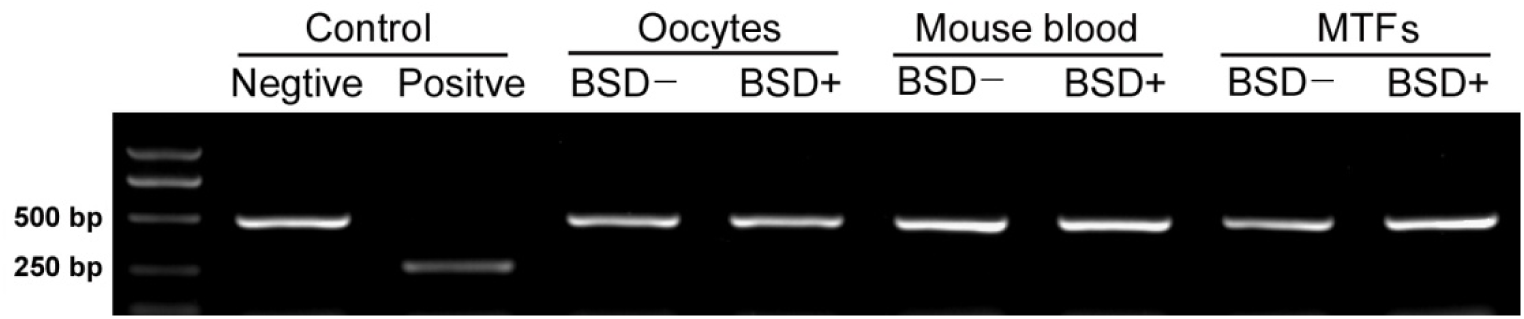
The electrophorogram of mycoplasma PCR detection. The oocytes, blood and MTFs from BSD+ and BSD- mice were examined. The mycoplasma negative sample should show a band at approximately 481 bp. The mycoplasma positive sample would show a band in the range of 270 ± 8 bp.

**Extended Data Figure 6.**
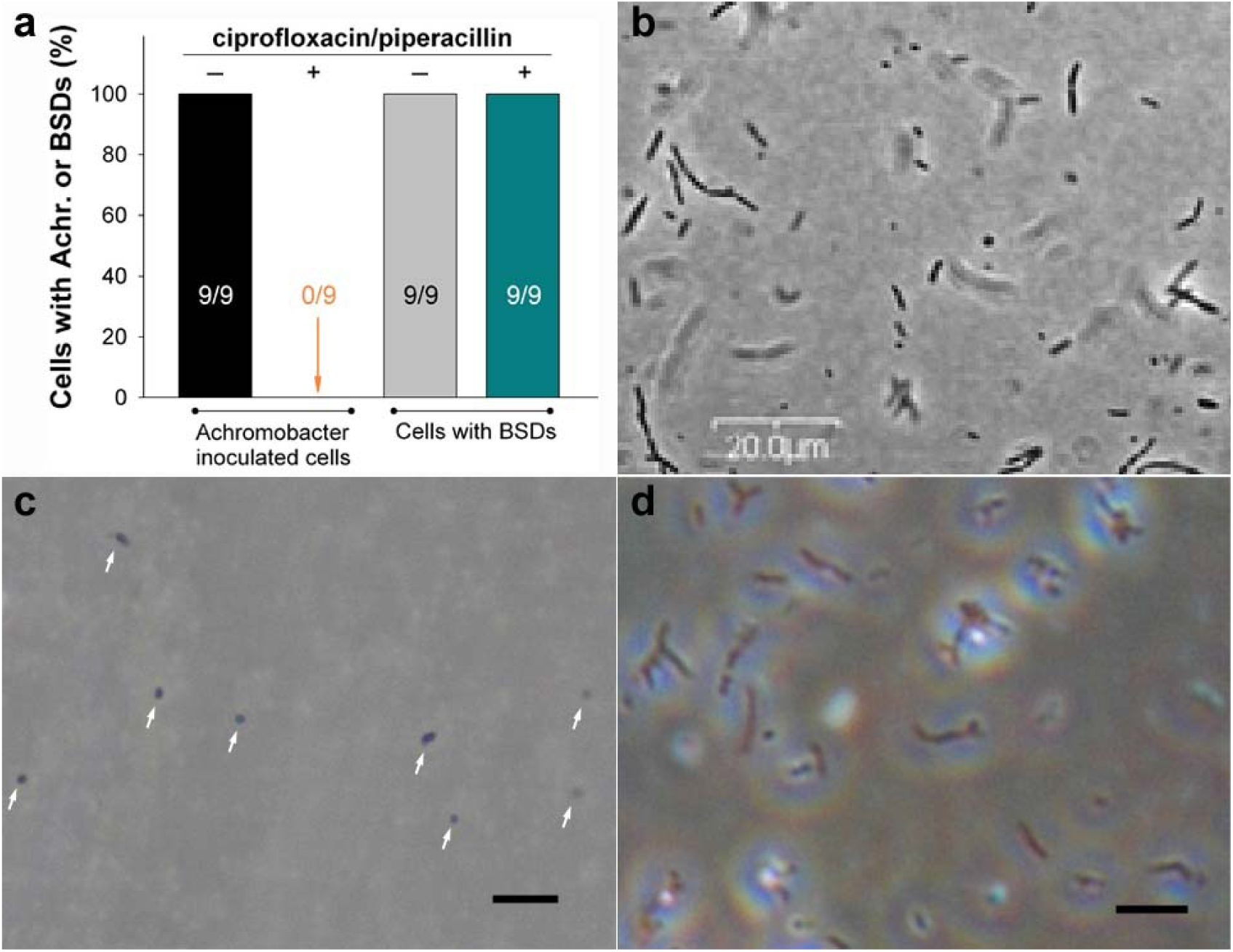
The difference in antibiotic resistance and morphology between BSDs and Achromobacter. **a**, The effects of ciprofloxacin/piperacillin on BSDs and Achromobacter. Each column denoted the percentage of cells with BSDs or Achromobacter. **b**, The image of Achromobacter contaminant published in Gray et al. paper ^5^. **c**, The morphology of BSDs (arrows) in cell dishes. Scale bar = 1 μm. **d**, The morphology of Achromobacter inoculated in BSD- cells. Scale bar = 10 μm. Comparison of **b, c, d** showing an obviously morphological difference between BSDs and Achromobacter.

**Extended Data Table 1.**
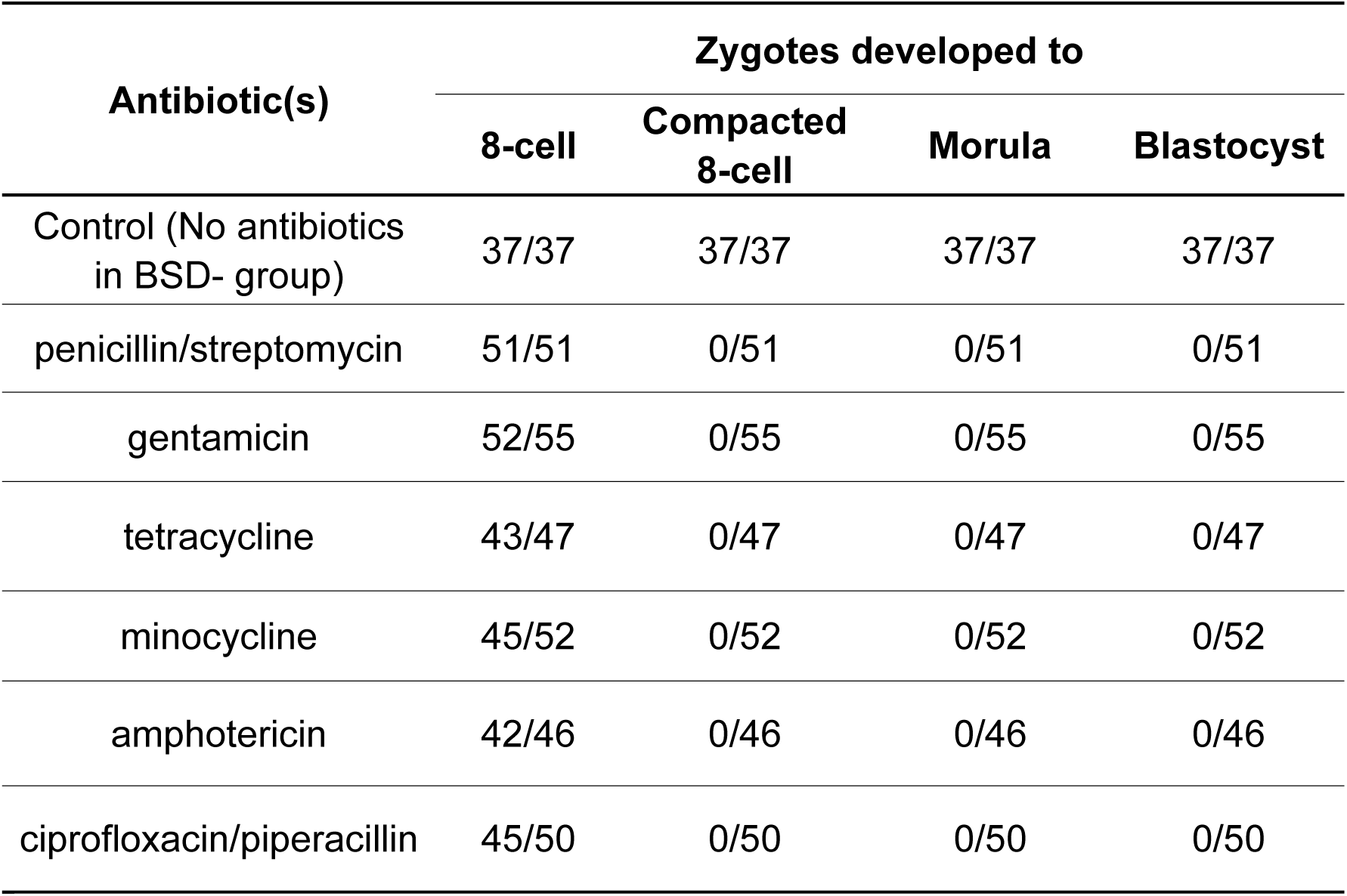
The effects of antibiotics on BSD+ embryo development *in vitro*.

| Antibiotic(s) | Zygotes developed to |  |  |  |
| --- | --- | --- | --- | --- |
|  | 8-cell | Compacted 8-cell | Morula | Blastocyst |
| Control (No antibiotics in BSD- group) | 37/37 | 37/37 | 37/37 | 37/37 |
| penicillin/streptomycin | 51/51 | 0/51 | 0/51 | 0/51 |
| gentamicin | 52/55 | 0/55 | 0/55 | 0/55 |
| tetracycline | 43/47 | 0/47 | 0/47 | 0/47 |
| minocycline | 45/52 | 0/52 | 0/52 | 0/52 |
| amphotericin | 42/46 | 0/46 | 0/46 | 0/46 |
| ciprofloxacin/piperacillin | 45/50 | 0/50 | 0/50 | 0/50 |

## Abbreviations

AAS: atomic absorption spectroscopy

BSDs: black swimming dots

EDX: energy dispersive X-Ray spectroscopy

HAP: hydroxyapatite

MEFs: mouse embryonic fibroblasts

MTFs: mouse tail-tip fibroblasts

NB: nanobacteria

SEM: scanning electronic microscope

TEM: transmission electronic microscope

## References

1 Kajander, E. O. & Ciftcioglu, N. Nanobacteria: an alternative mechanism for pathogenic intra- and extracellular calcification and stone formation. Proc. Natl. Acad. Sci. U. S. A. 95, 8274–8279 (1998).

2 Cisar, J. O. et al. An alternative interpretation of nanobacteria-induced biomineralization. Proc. Natl. Acad. Sci. U. S. A. 97, 11511–11515 (2000).

3 Martel, J. & Young, J. D. Purported nanobacteria in human blood as calcium carbonate nanoparticles. Proc. Natl. Acad. Sci. U. S. A. 105, 5549–5554 (2008).

4 Çiftçioǧlu, N., Haddad, R. S., Golden, D. C., Morrison, D. R. & Mckay, D. S. A potential cause for kidney stone formation during space flights: Enhanced growth of nanobacteria in microgravity. Kidney Int. 67, 483–491 (2005).

5 Gray, J. S., Birmingham, J. M. & Fenton, J. I. Got black swimming dots in your cell culture? Identification of Achromobacter as a novel cell culture contaminant. Biologicals 38, 273–277 (2010).

6 Folk, R. L. SEM imaging of bacteria and nannobacteria in carbonate sediments and rocks. J. Sediment. Res. 63, 990–999 (1993).

7 Akernan, K. K., Kuronen, I. & Kajander, E. O. Scanning Electron Microscopy of Nanobacteria -Novel Biofilm Producing Organisms in Blood. Scanning 15 (1993).

8 Martel, J., Peng, H.-H., Young, D., Wu, C.-Y. & Young, J. D. Of nanobacteria, nanoparticles, biofilms and their role in health and disease: facts, fancy and future. Nanomedicine 9, 483–499 (2014).

9 Young, J. D. & Martel, J. The rise and fall of nanobacteria. Sci. Am. 302, 52–59 (2010).

10 Dorozhkin, S. V. Nanodimensional and Nanocrystalline Apatites and Other Calcium Orthophosphates in Biomedical Engineering, Biology and Medicine. Materials 2, 1975–2045 (2009).

11 Dorozhkin, S. V. & Epple, M. Biological and Medical Significance of Calcium Phosphates. Angew. Chem. Int. Edit. 41, 3130–3146 (2002).

12 Urbano, P. & Urbano, F. Nanobacteria: facts or fancies? PLoS Pathog. 3, e55, doi:10.1371/journal.ppat.0030055 (2007).

13 Tian, C. Y., Xu, J. J. & Chen, H. Y. A novel aptasensor for the detection of adenosine in cancer cells by electrochemiluminescence of nitrogen doped TiO2 nanotubes. Chem. Commun. 48, 8234 (2012).

14 Laskin, A. & Cowin, J. P. Automated single-particle SEM/EDX analysis of submicrometer particles down to 0.1 microm. Anal. Chem. 73, 1023–1029 (2001).

15 Ciftcioglu, N. & McKay, D. S. Pathological Calcification and Replicating Calcifying-Nanoparticles:General Approach and Correlation. Pediatr. Res. 67, 490–499 (2010).

16 Chave, K. E. Physics and Chemistry of Biomineralization. Annu. Rev. Earth Pl. Sc. 12, 293–305 (2003).

17 Simkiss, K. Biomineralization and detoxification. Calcif. Tissue Res. 24, 199–200 (1977).

18 Bawden, J. W. Calcium transport during mineralization. Anat. Rec. 224, 226–233 (1989).

19 Wong, T.-Y. et al. Detection and characterization of mineralo-organic nanoparticles in human kidneys. Sci. Rep. 5, 15272 (2015).

## References

20 Takahashi, K. & Yamanaka, S. Induction of pluripotent stem cells from mouse embryonic and adult fibroblast cultures by defined factors. Cell 126, 663–676 (2006).

21 Wang, Y. et al. Modifications of chemically induced-enucleated nuclear transfer technique by reverse-order nuclear transfer in mouse. Zygote 17, 261–268 (2009).

